# Response to immune checkpoint blockade improved in pre-clinical model of breast cancer after bariatric surgery

**DOI:** 10.1101/2022.03.30.486293

**Authors:** Laura M. Sipe, Mehdi Chaib, Emily B. Korba, Heejoon Jo, Mary-Camille Lovely, Brittany R. Counts, Ubaid Tanveer, Jared C. Clements, Neena A. John, Deidre Daria, Tony N. Marion, Radhika Sekhri, Ajeeth K. Pingili, Bin Teng, James A. Carson, D. Neil Hayes, Matthew J. Davis, Joseph F. Pierre, Liza Makowski

## Abstract

Bariatric surgery is becoming more prevalent as a sustainable weight loss approach, with vertical sleeve gastrectomy (VSG) being the first line of surgical intervention. We and others have shown that obesity exacerbates tumor growth while diet-induced weight loss impairs obesity-driven progression. It remains unknown how bariatric surgery-induced weight loss impacts cancer progression or alters responses to therapy. Using a pre-clinical model of diet induced obesity followed by VSG or diet-induced weight loss, breast cancer progression and immune checkpoint blockade therapy was investigated. Weight loss by bariatric surgery or weight matched dietary intervention before tumor engraftment protected against obesity-exacerbated tumor progression. However, VSG was not as effective as dietary intervention in reducing tumor burden despite achieving a similar extent of weight and adiposity loss. Circulating leptin did not associate with changes in tumor burden. Uniquely, tumors in mice that received VSG displayed elevated inflammation and immune checkpoint ligand, PD-L1. Further, mice that received VSG had reduced tumor infiltrating T lymphocytes and cytolysis suggesting an ineffective anti-tumor microenvironment. VSG-associated elevation of PD-L1 prompted us to next investigate the efficacy of immune checkpoint blockade in lean, obese, and formerly obese mice that lost weight by VSG or weight matched controls. While obese mice were resistant to immune checkpoint blockade, anti-PD-L1 potently impaired tumor progression after VSG through improved anti-tumor immunity. Thus, in formerly obese mice, surgical weight loss followed by immunotherapy reduced breast cancer burden.

## Introduction

Obese breast cancer patients, defined as having a body mass index greater than 30, have worsened breast cancer prognoses with elevated breast cancer invasion [1, 2], distant metastases [3–5], tumor recurrence [6, 7], impaired delivery of systemic therapies [8, 9], and high mortality [10–12]. Weight loss interventions focusing on dietary approaches and exercise have demonstrated improved prognoses after a breast cancer diagnosis [13–17]. Pre-clinical models support that weight loss through diet or physical activity prior to tumor onset is beneficial to reduce obesity associated tumor progression [18–22]. Thus, intentional weight loss prior to tumor onset is a potential intervention to reduce negative cancer outcomes.

Bariatric surgery, also known as metabolic surgery, is an effective intervention for obese patients that leads to stable and sustained weight loss. Bariatric surgery primarily encompasses gastric banding, Roux-en-Y gastric bypass, and vertical sleeve gastrectomy (VSG) [23]. VSG is currently the least invasive and most common bariatric procedure [24]. Patients who receive a VSG have a reduction of 57% excess weight after two years, which remains relatively stable out to 10 years post surgery [25]. Remarkably, patients who undergo surgically induced weight loss have a reduction in all-cause mortality up to 60% [26, 27].

Here, to investigate the impacts of obesity and bariatric surgery-induced weight loss on breast cancer progression and response to therapy, we utilized female C57BL/6J mice, which are obesogenic and immune competent. Once obese, mice were subjected to weight loss interventions including bariatric surgery by VSG or dietary intervention as a weight matched control. Mice not subjected to VSG received a control sham surgery. Mice remaining obese or formerly obese mice that lost weight by surgery or diet were subsequently implanted orthotopically with syngeneic breast cancer cells to determine impacts on tumor progression, burden, and anti-tumor immunity. We found that mice that received the VSG displayed reduced obesity-accelerated breast cancer compared to obese sham treated controls. However, the most effective blunting of tumor progression was detected in weight matched sham controls. Thus, bariatric surgery was effective at reducing tumor burden but not to the same extent as weight matched controls despite similar weight and adiposity loss. A potential mediator of moderate impacts on tumor progression after VSG was the upregulation of checkpoint ligand, programmed death ligand 1 (PD-L1) and reduced CD8+ T cell content in tumors uniquely in VSG-treated mice. Thus, to determine if immune checkpoint blockade (ICB) after VSG could improve tumor outcomes, we report that in mice after VSG, anti-PD-L1 was efficacious to reduce breast cancer burden comparable to lean controls, while obese mice were resistant to anti-PD-L1. In sum, our study contributes critical observations regarding the impacts of obesity and bariatric surgery-induced weight loss on breast cancer progression and response to immunotherapy that are relevant to this rapidly emerging area of research and medicine.

## Results

### Surgical and dietary weight loss interventions reduced weight to the same extent

To quantify impacts of bariatric surgery on cancer progression, weight loss was induced prior to tumor implantation (study design, **Fig. 1A**). Female C57BL/6J mice were weaned onto low fat diet (LFD) to remain lean or onto high fat diet (HFD) to become obese. After 16 weeks on diet, HFD-fed mice displayed marked diet induced obesity (DIO, **Fig. 1B**). A subset of DIO mice then underwent surgical or dietary weight loss interventions. Surgically treated DIO mice received the VSG bariatric procedure, wherein the lateral 80% of the stomach was removed and the remaining stomach was sutured creating a tubular gastric sleeve [28]. VSG induced a significant and sustained weight loss of 20% of the starting body weight, despite being continuously maintained on HFD (HFD-VSG, **Fig. 1C,** detailed statistical comparisons within **Table S1**). HFD-VSG mice lost weight to within a few grams of lean LFD-sham treated control mice. Importantly, mice did not regain weight after the VSG. Weight rebound has often been recorded in other studies in this time course [29, 30]. To control for the effects of surgery, all other groups that did not undergo a VSG received a sham surgery including perioperative procedures, abdominal laparotomy, anesthesia, and analgesics with minimal impacts on weight maintenance (**Fig. 1A, 1C**). To compare the impact of VSG on breast cancer outcomes to weight loss *per se*, we employed a dietary weight loss intervention initiated after sham surgery wherein mice were fed calorically restricted amounts of HFD to match the weight loss and diet exposure of HFD-VSG treated mice, termed weight matched sham (WM-Sham). As designed, WM-Sham body weight loss was not significantly different from HFD-VSG (**Fig. 1C**). By endpoint, five weeks after surgical and diet interventions, both weight loss groups (HFD-VSG and WM-Sham) displayed significantly reduced body weights compared to HFD-Sham obese control mice (**Fig. 1C**). These results demonstrate successful generation of complementary weight loss approaches to next investigate the impacts of bariatric surgery-mediated weight loss on tumor progression.

**Figure 1.**
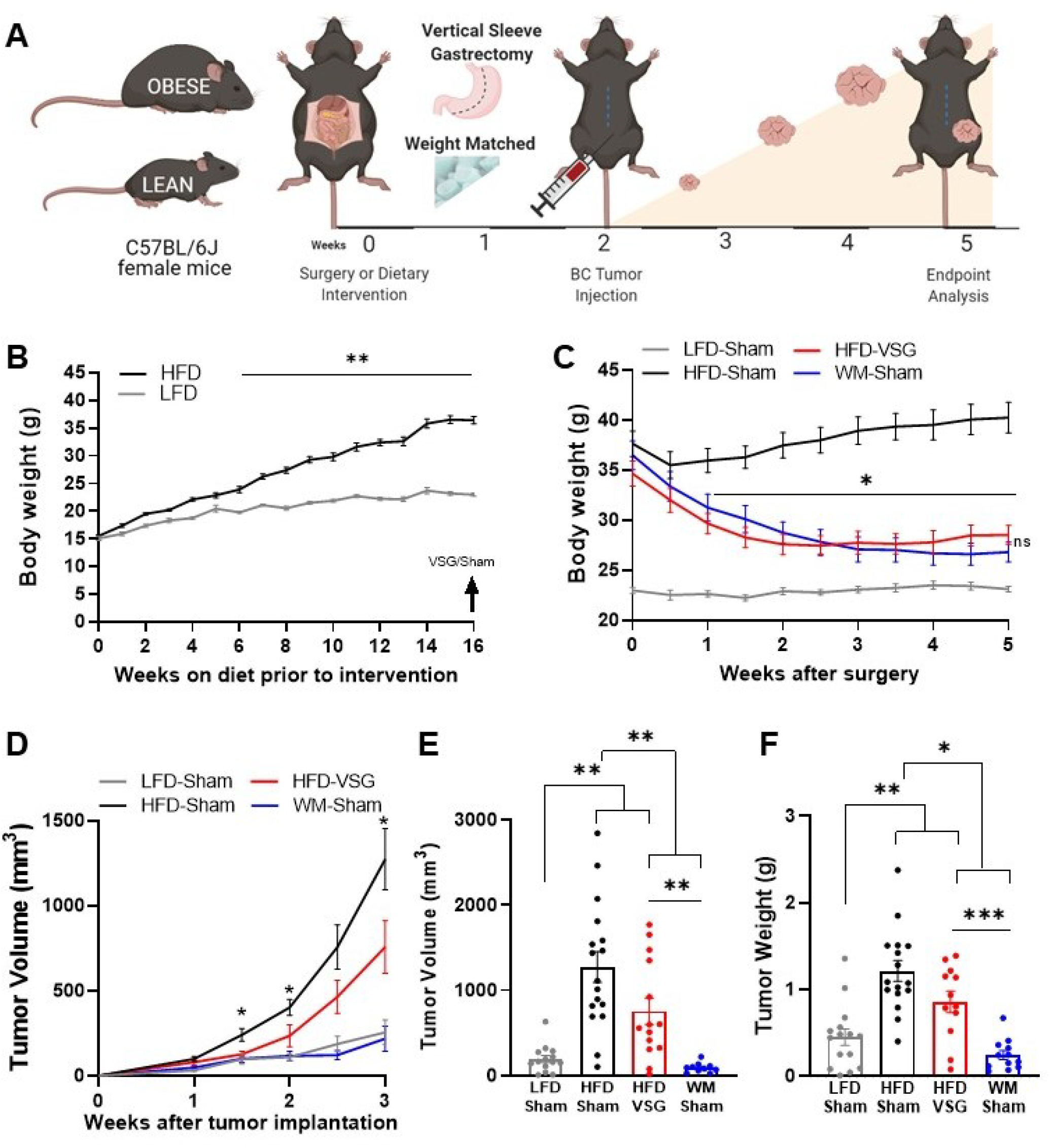
Surgical and dietary weight loss interventions reduced tumor progression and burden compared to obese mice. (**A**) Schematic of diet induced obesity, weight loss intervention, and breast cancer cell injection in female C57BL/6J mice. Mice were fed obesogenic diets or kept lean for 16 weeks. At 20 weeks of age mice were subjected to bariatric surgery or dietary intervention and sham surgery to stably reduce weights while control high fat diet (HFD) and low fat diet (LFD) fed mice received sham surgery to remain obese or lean, respectively. E0771 breast cancer cells were injected at 22 weeks of age when weight loss stabilized. Tumor progression was quantified, and mice were sacrificed at endpoint 3 weeks later. (**B**) Weekly body weights are shown as DIO is established over 16 weeks on HFD compared to lean control mice fed LFD (n=15). (**C**) Body weights were measured biweekly after DIO mice were subjected to either bariatric surgery or dietary weight loss interventions. Four groups include: HFD-fed and vertical sleeve gastrectomy (HFD-VSG, red) and weight-matched (WM) caloric restricted HFD-fed and sham (WM-Sham, blue) to mirror weight loss in VSG group. These interventions were compared to controls continuously HFD-fed and sham (HFD-Sham, black) or continuously LFD-fed and sham (LFD-Sham, grey). (**D**) Tumor volume quantified over three weeks. (C-D) Two-way ANOVA Fisher’s LSD test for individual comparisons with *p<0.05, **p<0.01 signifying HFD-Sham compared to all other groups and detailed in table S1 and S2 respectively(**E**) Tumor volume and (**F**) tumor weight at endpoint. (E-F) Mean ± SEM One-way ANOVA with Fisher’s LSD test. (B-F) n=15 LFD-Sham, n=17 HFD-Sham, n=14 HFD-VSG, n=13 WM-Sham. Mean ± SEM *p<0.05, **p<0.01, ***p<0.001.

### Obesity-accelerated breast cancer progression was reversed by VSG and dietary weight loss

To determine if surgical weight loss corrects obesity-associated breast cancer progression, E0771 syngeneic breast cancer cells were orthotopically implanted into the 4^th^ mammary fat pad two weeks following weight loss interventions, when weight loss was stabilized (**Fig. 1A, 1C**). Tumor progression was quantified over 3 weeks (**Fig. 1A, 1D,** detailed statistics within **Table S2**). Breast cancer cell implantation and progression did not adversely impact body weight (**Fig. 1C**). HFD-Sham tumors were significantly larger than LFD-Sham by 1 week after cell implantation. In mice that had lost weight, reduced tumor progression was observed compared to HFD-Sham from 1.5 weeks after implantation (**Fig. 1D**). At endpoint, HFD-VSG tumors were significantly smaller than HFD-Sham by volume and weight (**Fig.1 D-F**). However, tumors in the WM-Sham group were significantly smaller than HFD-VSG despite identical body weights between the two weight loss approaches (**Fig. 1C-F**). In fact, tumor progression was blunted in WM-Sham controls such that at endpoint tumors in WM-Sham were not significantly different from tumors in LFD-Sham lean controls by volume or weight (**Fig. 1D-F**). Thus, dietary intervention in formerly obese mice was most impactful to restore a lean-like tumor phenotype with minimal tumor progression evident and the smallest tumor burden, while weight loss by VSG proved to be less impactful to blunt tumor progression compared to weight matched controls.

### Adiposity and leptin were reduced in formerly obese mice

Increased adiposity is associated with obesity-worsened breast cancer [31]. Surgical and dietary interventions resulted in a significant reduction in adiposity compared to HFD-Sham obese control mice as early as week one post-surgery that stabilized two weeks after intervention and persisted until endpoint (**Fig. 2A**). Breast cancer cell implantation and progression from weeks 2-5 did not impact adiposity in any group (**Fig. 2A**). In line with adiposity, HFD-Sham mice had about 10-fold greater mammary fat pad and gonadal adipose mass compared to lean LFD-Sham controls (**Fig. 2B-C**). HFD-VSG and WM-Sham groups lost significant adipose mass compared to HFD-Sham obese controls, but not to the extent quantified in lean LFD-Sham mice (**Fig. 2A-C**). Enlarged adipocyte size in the mammary fat pad is a mediator of obesity associated inflammation and impacts breast cancer progression [32]. Adipocyte size in the mammary fat pad was enlarged in HFD-Sham compared to LFD-Sham mice (**Fig. 2D**). HFD-VSG mammary fat pads contained significantly smaller adipocytes compared to HFD-Sham but did not reduce size to that of LFD-Sham (**Fig. 2D**). Interestingly, WM-Sham mice retained significantly larger adipocytes compared to HFD-VSG, despite similar loss of adiposity and identical mammary fat pad and gonadal adipose depot weights (**Fig. 2A-D**). Therefore, the association with greater adipocyte size and larger tumor burden did not hold true in these models of formerly obese mice.

**Figure 2.**
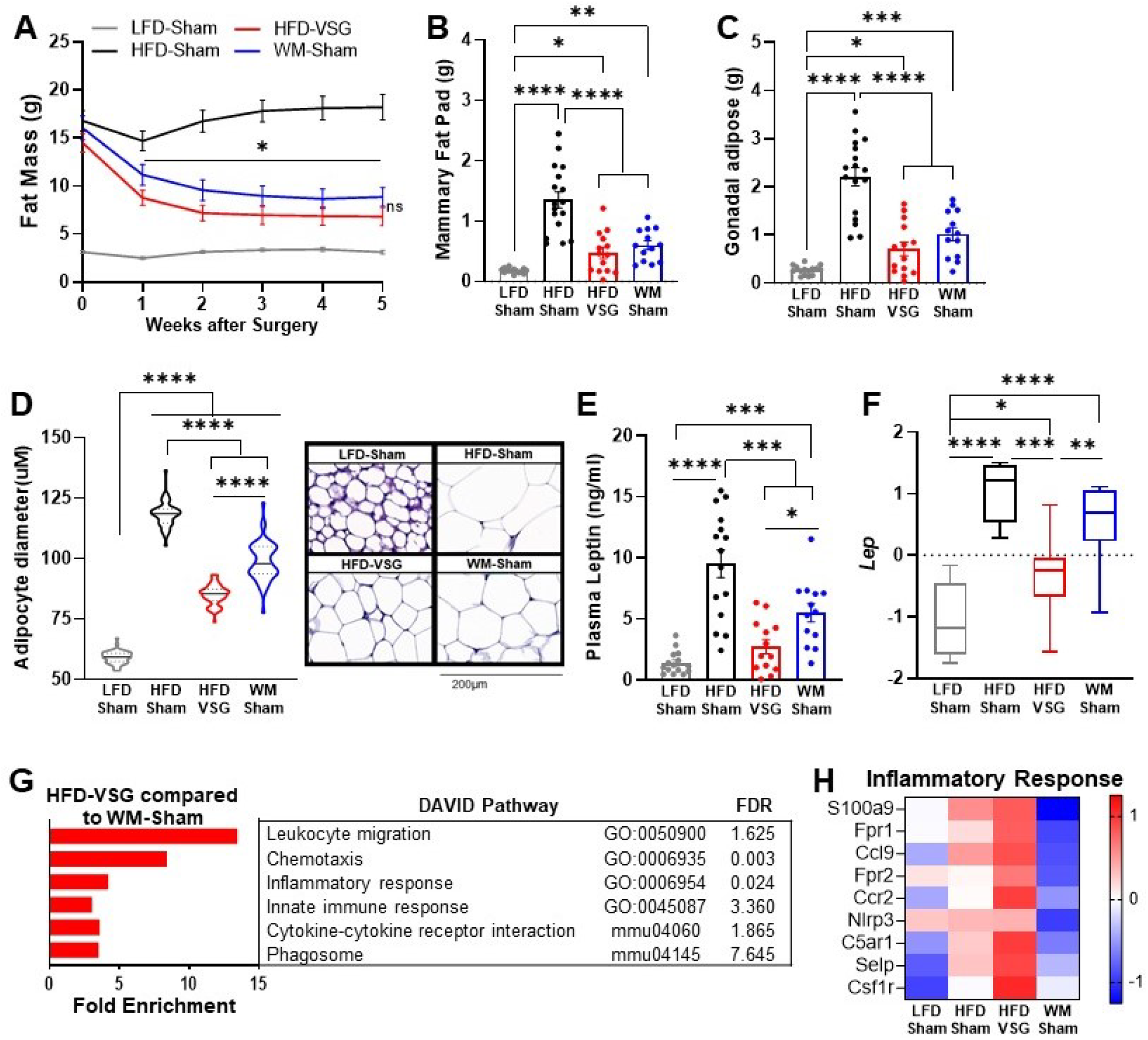
Bariatric surgery reduced adiposity similarly to weight matched controls, yet increased inflammation in mammary fat pad. (**A**) Fat mass was measured by EchoMRI. Mean ± SEM is shown. Two-way ANOVA with Fisher’s LSD Test, *p<0.05 all other groups compared to HFD-Sham. (**B**) Mammary fat pad and (**C**) gonadal adipose weights were measured at endpoint. (A-C) Mean ± SEM is shown. n=15 LFD-Sham, n=17 HFD-Sham, n=14 HFD-VSG, n=13 WM-Sham. (**D**) Adipocyte diameter along the longest length was measured in H&E sections of uninjected contralateral mammary fat pad. Violin plot with median (solid line) and quartiles (dashed line) is shown. Representative images at 20X are shown with 200μm represented by scale bar. N=5-7, n=50 adipocytes/sample. (**E**) Leptin concentration in plasma was measured at endpoint after 4 hours of fasting. N=13-15. (**F**) Row mean centered gene expression of *Lep* encoding for Leptin in uninjected contralateral mammary fat pad by RNA-seq. Box and whiskers shown mean, min, and max. N=6-8. (B-E). One way ANOVA with Fisher’s LSD test. *p<0.05, **p<0.01, ***p<0.001, ****p<0.0001. (**G**) DAVID analysis of regulated inflammatory pathways in mammary fat pads of HFD-VSG mice compared to WM-Sham mice. FDR, false discovery rate. (**H**) Heat map of row mean centered gene expression in uninjected contralateral mammary fat pad by RNA-seq of genes contributing to the significantly regulated Inflammatory Response Pathway (GO:0006954) determined by DAVID analysis. N=6-8.

Leptin is associated with adiposity and adipocyte size and can signal to activate breast cancer cell proliferation [33]. Plasma leptin concentrations (**Fig. 2E**) and leptin mRNA expression in mammary fat pad (**Fig. 2F**) paralleled findings for endpoint adipocyte size (**Fig. 2D**) with HFD-Sham displaying the greatest leptin plasma concentrations and mammary fat pad expression. HFD-VSG reduced leptin concentrations in plasma and in adipocytes compared to HFD-Sham obese controls (**Fig. 2E-F**). As in adipocyte size, despite comparable weight loss and adipose mass between VSG and WM-Sham groups, WM-Sham had 2-fold greater leptin concentration in plasma or expression in mammary fat pad compared to HFD-VSG (**Fig. 2E-F**). Thus, leptin mediated signaling does not account for why VSG is less effective in reducing tumor burden compared to weight loss alone.

### Elevated inflammation was evident in mammary fat pad uniquely after VSG weight loss intervention

Increased inflammation in the adipose has been reported in mouse models of VSG, with persistent elevations in adipose tissue macrophages despite improvements in obesity-associated parameters [34–37]. Thus, we investigated if inflammatory changes in the mammary fat pad reflect pathways that could impact tumor burden [33]. Compared to WM-Sham controls, HFD-VSG mammary fat pads reflected 5-10-fold elevation of immune pathways such as leukocyte migration, chemotaxis, inflammatory response, among others (**Fig. 2G**). Examining key genes common to the inflammatory response pathways, compared to LFD-Sham lean controls, HFD-Sham obese mice displayed elevated expression of many inflammatory genes such as chemokine receptor *Ccr2* and growth factor receptor *Csf1r*, among others, as expected with DIO (**Fig. 2H**). Despite significant reductions in adiposity and adipocyte size after VSG, mammary fat pads from HFD-VSG mice displayed evidence of persistent or exacerbated inflammation compared to all groups including HFD-Sham obese controls (**Fig. 2H**). In stark contrast, compared to both HFD-Sham and HFD-VSG groups, mammary fat pads from WM-Sham treated mice displayed greatly reduced inflammatory gene expression to levels similar to, or lower than, lean LFD-Sham controls (**Fig. 2H**). Taken together, the increased inflammatory response signature in the mammary fat pads of HFD-VSG mice suggests the possibility of a more tumor permissive environment, particularly compared to WM-Sham controls.

### Tumors displayed elevated inflammation and immune checkpoint ligand expression in mice receiving VSG

Like the mammary fat pad, transcriptome analysis of tumors in mice after VSG intervention displayed increased enrichment of inflammatory response as well as response to hypoxia pathways compared to HFD-Sham tumors, indicating an inflamed and hypoxic tumor microenvironment (**Fig. 3A**), whereas these pathways were downregulated in tumors from WM-Sham mice (**Fig. 3A**). Elevated pathways in VSG tumors (**Fig. 3A**) contain genes - specifically *Tlr2, Tlr13, Ifngr1, Ccl9, Hif1a*, and *Cybb* - that are established to increase immune checkpoint ligand PD-L1 expression (**Fig. 3B**) [38, 39]. Therefore, we next queried immune checkpoint expression in the tumor microenvironment to determine if elevated pathways and genes in the VSG-treated group could lead to increased immune checkpoint ligand expression. Indeed, flow cytometry analysis revealed that the frequency of PD-L1 + cells was significantly and uniquely elevated in tumors after VSG intervention compared to all other groups in the CD45-fraction (**Fig. 3C**). The CD45-fraction contains tumor cells as well as other stromal cells such as fibroblasts, endothelial cells, adipose stromal cells etc. Furthermore, expression of PD-L1 quantified by MFI was also significantly elevated in the CD45-fraction from HFD-VSG tumors (**Fig. 3D**). In contrast, WM-Sham intervention significantly reduced frequency of PD-L1+ non-immune cells and PD-L1 MFI relative to tumors from HFD-VSG treated mice by 60 and 30%, respectively (**Fig. 3C-D**). Overall, surgically induced weight loss increased tumor inflammation and elevated the immune checkpoint ligand PD-L1 in the tumor microenvironment suggesting the presence of impaired anti-tumor immunity [40, 41].

**Figure 3.**
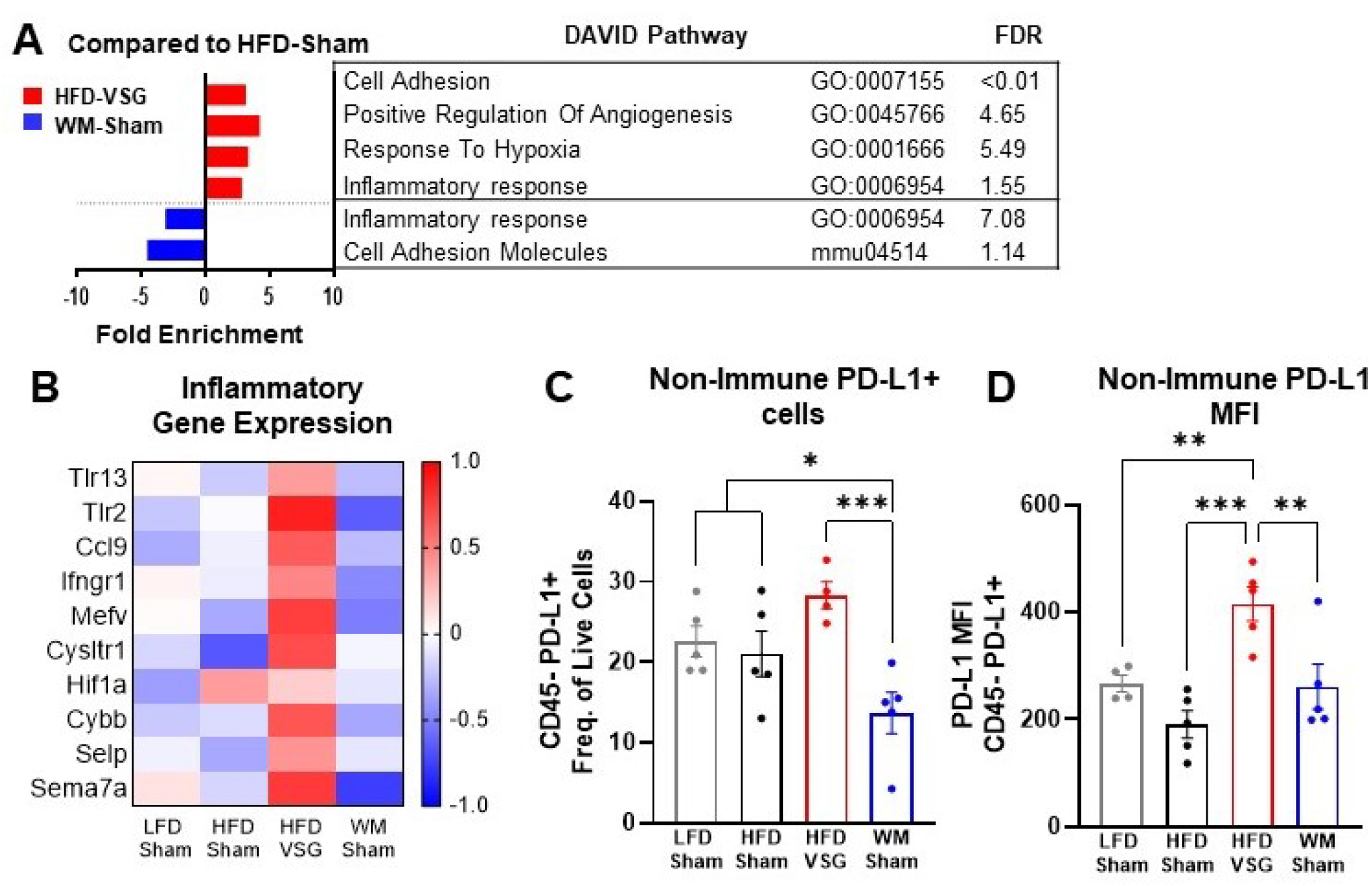
The tumor microenvironment displayed increased inflammation and immune checkpoint ligand expression following bariatric surgery. (**A**) DAVID analysis of regulated pathways and false discovery rate (FDR) for HFD-VSG (red) and WM-Sham (blue) relative to tumors from HFD-Sham mice is shown. N= 6-8. (**B**) Heat map of row mean centered gene expression in tumor by RNA-seq of genes contributing to significantly regulated inflammatory response pathway (GO:0006954) and response to hypoxia pathway (GO:0001666) determined by DAVID analysis. N=6-8. (**C**) Flow cytometric analysis of CD45 negative (CD45-) PD-L1+ non-immune cells in tumor are plotted as frequency of total live cells. (**D**) Mean fluorescent intensity (MFI) of PD-L1 on CD45-PD-L1+ cells in tumor are shown. N=4-5. Mean ± SEM is shown. One-way ANOVA with Fisher’s LSD test. *p<0.05, **p<0.01, ***p<0.001.

### T cell tumor infiltration and cytolysis were impaired after VSG

In the tumor microenvironment, high PD-L1 expression by tumor cells can dampen T cell-mediated anti-tumor immune responses [40, 41]. Therefore, we next investigated T cell content and associated activation pathways [42]. CD3+ T cell frequency in tumors from HFD-VSG mice was significantly decreased compared to tumors from LFD-Sham control mice (**Fig. 4A**). In contrast, CD3+ T cell frequency in weight matched controls was significantly greater compared to content in tumors after VSG (**Fig. 4A**). Obesity has been shown to decrease CD8+ cytotoxic tumor infiltrating T cells [42, 43] which was evident, but not significant, in this study comparing lean LFD-Sham to obese HFD-Sham controls (**Fig. 4B-C**). Obesity-driven CD8+ T cell reductions were not corrected in tumors from formerly obese HFD-VSG mice by both flow and RNA-seq CIBERSORT analysis (**Fig. 4B-C**). Importantly, obesity-driven reductions in CD8+ T cell frequencies were reversed in tumors from WM-Sham control mice and corrected to levels found in tumors from lean LFD-Sham controls (**Fig. 4B-C**). Transcriptomic analysis revealed that T cell specific signaling pathways and genes in the tumor mirrored T cell infiltration (**Fig. 4D-E**). Lowest T cell signaling gene signature expression was evident in tumors from HFD-Sham and HFD-VSG mice, with some correction in WM-Sham mice towards levels detected in lean LFD-Sham controls (**Fig. 4D-E**). Of note, CD3+ and CD8+ T cell frequencies were unchanged in the tumor adjacent mammary fat pad and tumor draining lymph node (TdLN) (**Fig. S1A-B**), suggesting T cell changes were specific to the tumor microenvironment. Further, neither T cells in tumor nor TdLN displayed changes in PD-1 expression measured by MFI (**Fig. S1C-D**).

**Figure 4.**
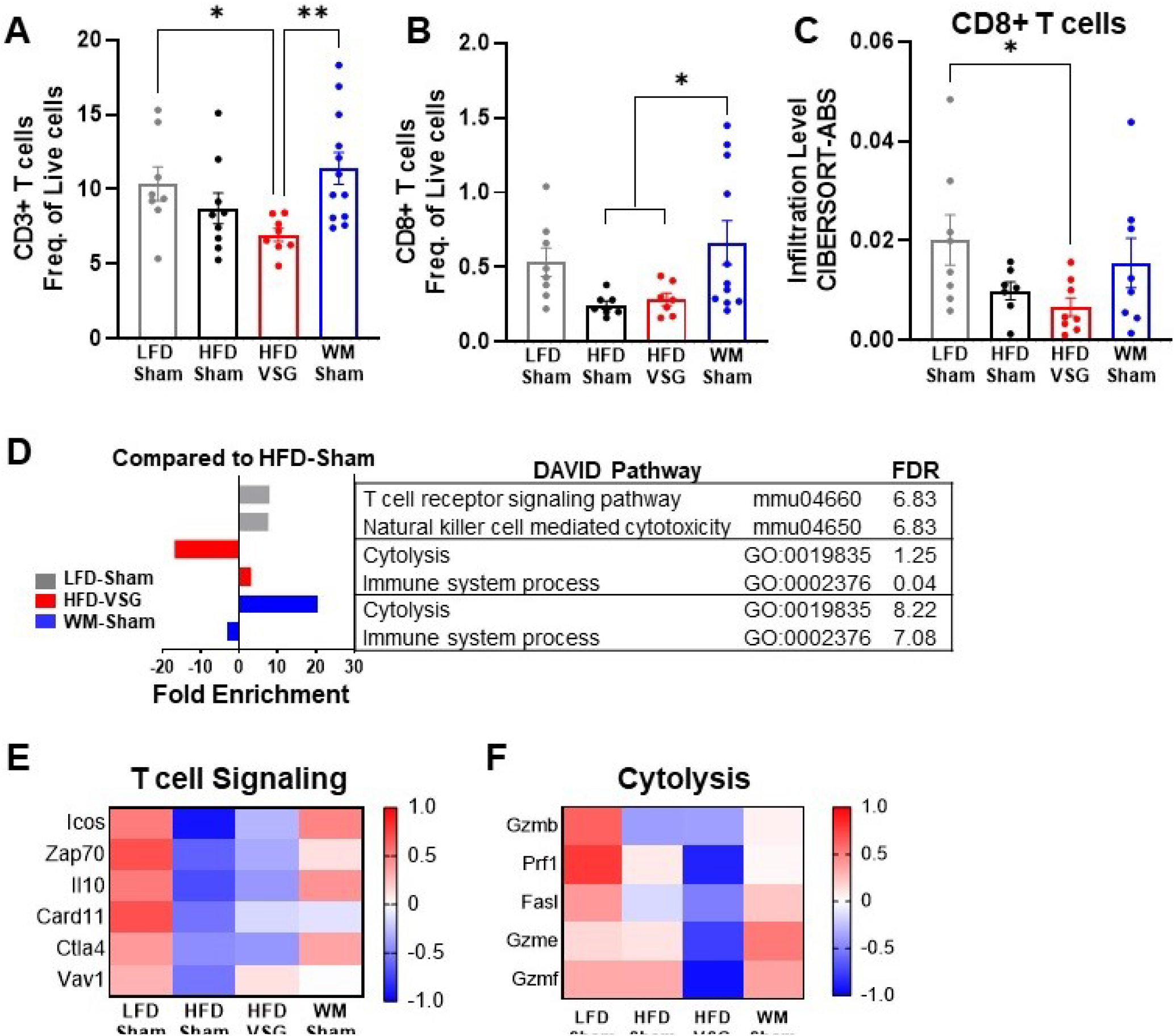
VSG reduced CD8+ tumor T lymphocyte frequency and markers of T cell activation demonstrating impaired anti-tumor immunity. (**A-B**) Flow cytometric analysis of tumor (A) CD3+ T cells and (B) CD8+ T cells are shown as frequency of total live cells. N=8-12. (**C**) Analysis of tumor CD8+ T cell infiltration from RNA-seq data using the CIBERSORT-Abs algorithm in TIMER2.0. N=6-8. (A-C) Mean ± SEM One-way ANOVA with Fisher’s LSD test *p<0.05, **p<0.01, ***p<0.001, ****p<0.0001. (**D**) DAVID analysis of regulated pathways for LFD-Sham (grey), HFD-VSG (red), and WM-Sham (blue) relative to tumors from HFD-Sham mice. N= 6-8. (**E**) Heat map of row mean centered gene expression in tumor by RNA-seq of genes contributing to the significantly regulated T cell signaling pathway (mmu04660, FDR 6.83) and (**F**) Cytolysis (GO:0019835, FDR 1.25) as determined by DAVID analysis. N=6-8.

A critical function of anti-tumor immune cells is effective cytolytic activity [42]. RNA-seq analysis showed that the cytolysis pathway was significantly and potently downregulated by 17-fold in HFD-VSG tumors compared to obese HFD-Sham controls (**Fig. 4D**). In contrast, tumors from the WM-Sham intervention group displayed the greatest activation with over 20-fold increase in the cytolysis pathway (**Fig. 4D**). Genes in the cytolytic pathway were greatly downregulated in HFD-VSG tumors compared to all other groups including granzymes and fas ligand (*Gzmb, Prf1, Fasl, Gzme,* and *Gzmf*), while gene expression was reversed to lean-like levels in tumors from WM-Sham mice (**Fig. 4F**) Taken together, weight matched control mice displayed uniquely restored T cell content and signaling pathways that were depressed by obesity which suggests an apparent effective anti-tumor response aligning with reduced tumor burden. In contrast, mice after VSG displayed a tumor microenvironment that resembled persistent obesity, with reduced T cell content and cytolytic markers, despite comparable weight loss with weight matched controls.

### Anti-PD-L1 therapy was more efficacious in VSG mice

The elevation of tumor immune checkpoint ligand PD-L1 after bariatric surgery may be one mechanism that underlies why surgical weight loss was less effective in reducing obesity-worsened tumor growth compared to weight loss alone. Therefore, we hypothesized that ICB would re-invigorate the anti-tumor immune response in mice after VSG to reduce tumor burden. Mice were weaned onto diets and received surgical or dietary weight loss interventions prior to tumor engraftment as above (**Fig. 1A**). Mice were then treated with anti-PD-L1 or isotype control IgG2b. Anti-PD-L1 did not affect body weight, mammary fat pad, or gonadal adipose weight suggesting no negative impacts on systemic homeostasis (**Fig. S2A-C**). In LFD-Sham lean controls, despite the tumor being 6-fold smaller than in obese mice at baseline, anti-PD-L1 significantly reduced tumor growth over time (**Fig 5A**). HFD-Sham mice were completely resistant to ICB (**Fig 5A-B**). Notably, anti-PD-L1 significantly reduced tumor progression in HFD-VSG (**Fig. 5A**), with significantly reduced tumor volume at endpoint (**Fig. 5B**). In line with an already active anti-tumor immune response, ICB was moderately and insignificantly effective in WM-Sham mice (**Fig. 5A-B**). Thus, ICB was efficacious in reducing tumor progression in mice after HFD-VSG to sizes comparable to tumors found in lean mice.

**Figure 5.**
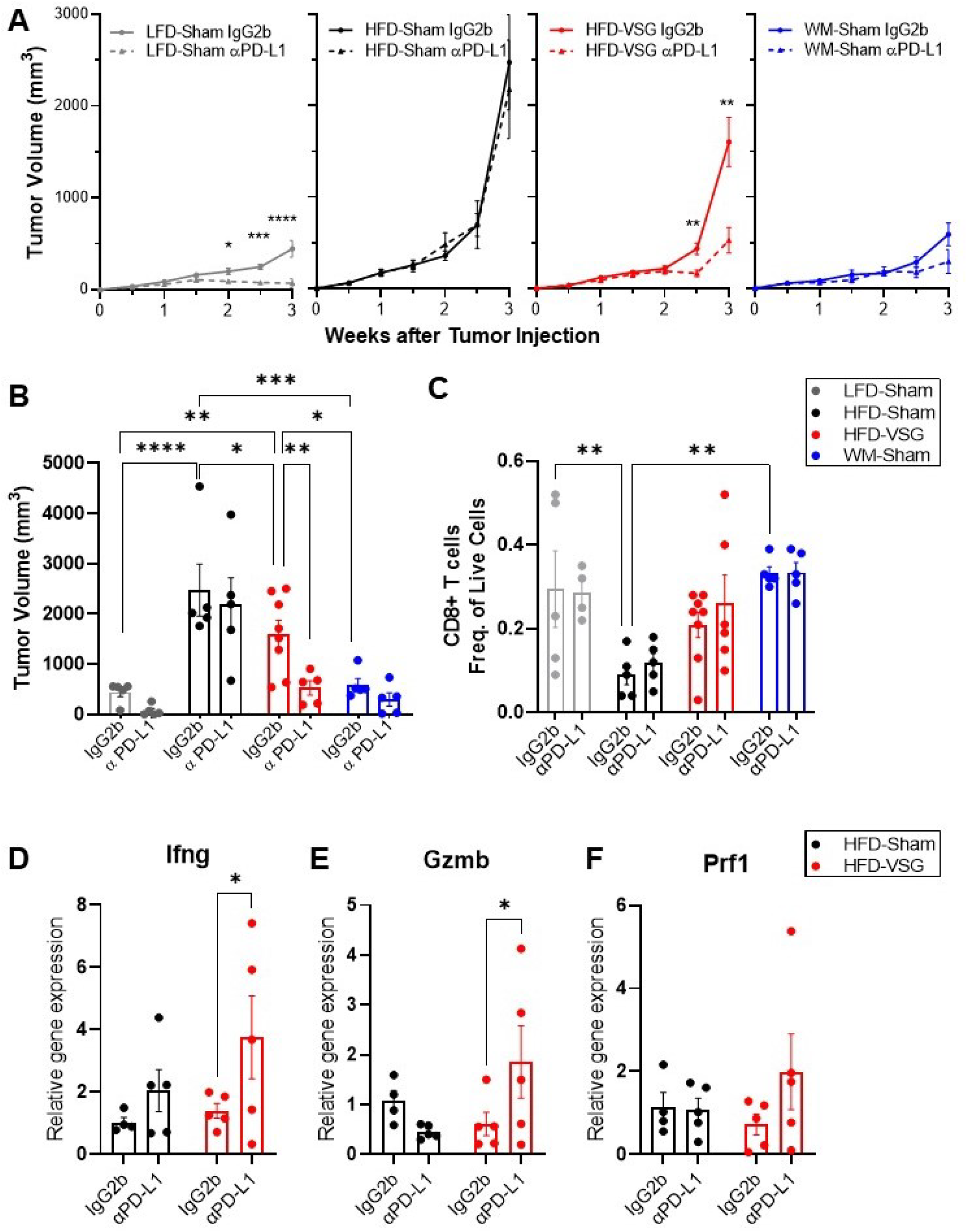
Immune checkpoint blockade reinvigorated the anti-tumor immune response in mice after bariatric surgery. DIO mice were subjected to either surgical or dietary weight loss interventions and compared to lean or obese controls similar to Fig 1A. After weight stabilization at 2 weeks, mice were injected with E0771 cells, as above. Mice were either treated with anti-PD-L1 or IgG2b isotype control every three days until sacrifice at 3 weeks after cell injection. (**A**) Mean tumor growth in each diet group treated with anti-PD-L1 or IgG2b isotype control is shown. (**B**) Tumor volume at endpoint. (**C**) Flow cytometric analysis of CD8+ T cells as frequency of total live cells in tumor. (**D**) Relative gene expression normalized to 18S of *Ifng* (**E**) *Gzmb* and (**F**) *Prf1* in tumors. (A-F) Mean ± SEM. N=5-8. Two-way ANOVA with Fisher’s LSD test. Only relevant statistical comparisons are shown for clarity. *p<0.05, **p<0.01, ***p<0.001, ****p<0.0001.

ICB restores cytotoxic T cell function, thus reestablishing effective anti-tumor immunity [40]. While there were not significant differences in mean CD8+ T cell content at endpoint (**Fig. 5C**), evidence of cytolytic capacity is upregulated in VSG tumors treated with anti-PD-L1 with increased *Ifng, Gzmb*, and *Prf1* expression (**Fig. 5D-F**). Our results suggest that ICB compensates for an ineffective anti-tumor immunity associated with elevated PD-L1 expression in the tumors of VSG mice to restore markers of cytotoxic T-cell response, which leads to reduced tumor burden.

### A <u>b</u>ariatric <u>s</u>urgery <u>a</u>ssociated weight loss <u>s</u>ignature derived from patient and murine adipose tissue associates with tumor burden

To determine if genes associated with weight loss after bariatric surgery are conserved across species, we compared subcutaneous adipose tissue biopsies from female human subject samples before and after bariatric surgery [37] with mammary fat pad tissue isolated from HFD-Sham and HFD-VSG mice in study 1 above (**Fig. 6A**). When comparing transcriptomic changes in adipose tissue after bariatric surgery from both humans and mouse models, there were 54 differentially expressed genes (DEGs) in common (**Fig. 6A**), which we termed the <u>B</u>ariatric <u>S</u>urgery <u>A</u>ssociated weight loss <u>S</u>ignature (BSAS, **Table S3**). Overlapping DEGs identified pathways involved in metabolism and adipose tissue remodeling after weight loss, and immune system processes (**Fig. 6B**). We next examined the relationship between BSAS and tumor burden in our model with divergent tumor growth patterns. Of the 54 genes in this BSAS, 11 genes significantly correlated to volumes of HFD-Sham and HFD-VSG tumors, which is shown in **Fig. 6C.** We termed these 11 genes the <u>T</u>umor associated BSAS (T-BSAS) gene signature (**Fig. 6C**). Seven of the genes were downregulated by obesity and reversed by VSG specific weight loss including *Ido1, Aldoc, Tmem125, Dgki, Slc7a4, Msc, and Ephb3*, while 4 were inversely regulated with obesity elevating *Klhl5, Nek6, Arhgap20*, and *Hp*. For example, *Ido1* expression in each group relative to tumor size is presented (**Fig. 6D**). Compared to the HFD-Sham obese group, the T-BSAS signature in HFD-VSG tumor largely resembled tumors from LFD-Sham (**Fig. 6D**). This multi-species approach uniquely demonstrates conserved transcriptional responses impacted by bariatric surgery that associate with tumor burden.

**Figure 6.**
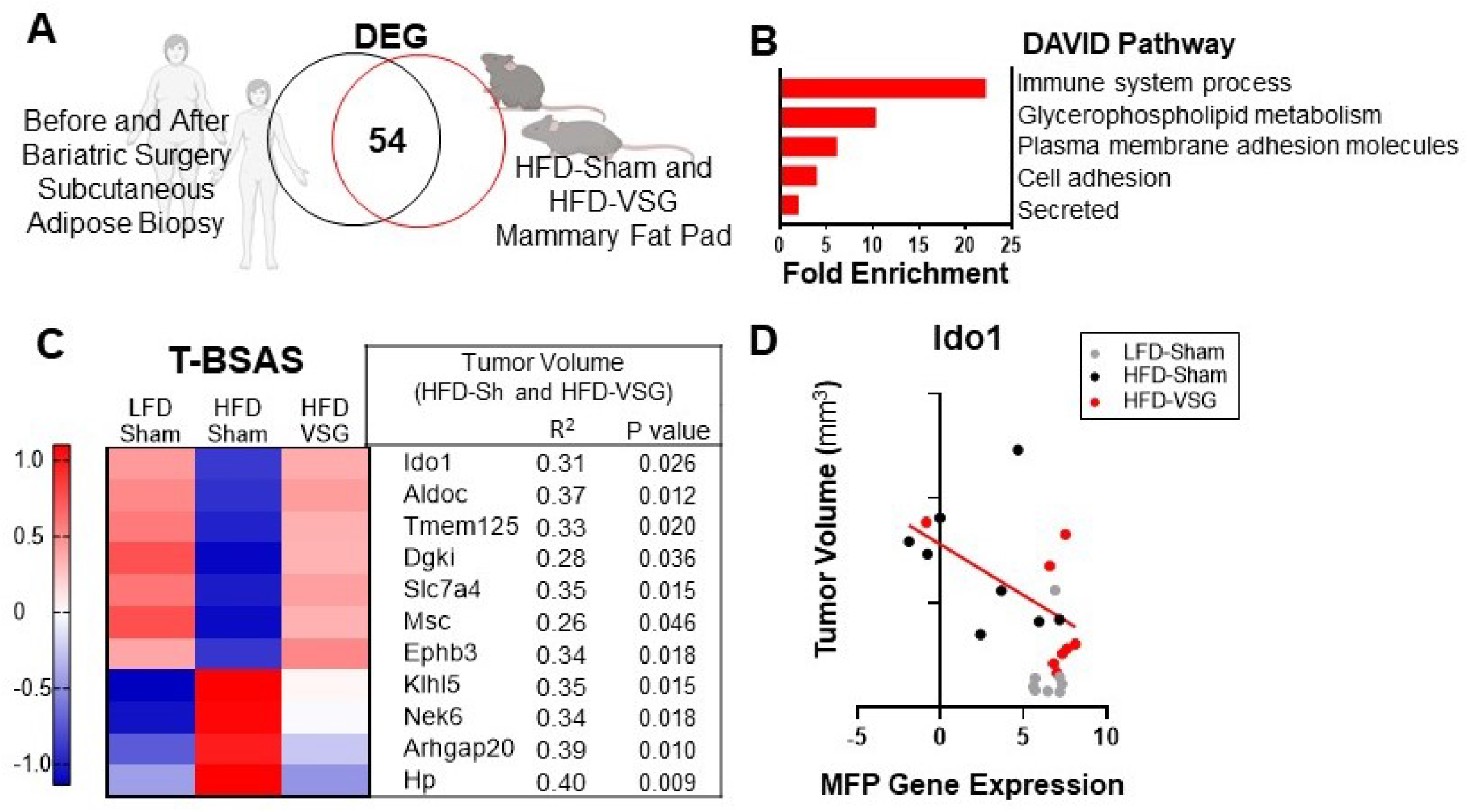
Conserved adipose bariatric surgery associated weight loss signature associated with tumor volume. (**A**) Venn diagram of differentially expressed genes (DEG) from obese and lean patient subcutaneous adipose tissue before and three months after bariatric surgery, respectively, compared to obese HFD-Sham and lean HFD-VSG mammary fat pad. (**B**) DAVID pathways enriched in the overlapping DEG are indicated. (**C**) A Tumor-<u>B</u>ariatric <u>S</u>urgery <u>A</u>ssociated weight loss <u>S</u>ignature (T-BSAS) signature was identified as a subset of BSAS genes that significantly correlated to tumor volume. Heat map of row mean centered expression of T-BSAS genes in the mammary fat pad by RNA-seq. (**D**) Tumor volume compared to unaffected mammary fat pad (MFP) gene expression of *Ido1* is plotted. Simple linear regression (red line) for HFD-Sham and HFD-VSG groups is shown (R^2^=0.31 and p=0.026).

## Discussion

Obesity negatively impacts many cancer outcomes and is thus a potential modifiable factor [23, 44]. In this study, to investigate the impacts of weight loss by bariatric surgery on subsequent tumor burden, we first established a murine model wherein once weight loss is stabilized, cancer cells were orthotopically implanted to examine progression and burden. We show that tumor growth in formerly obese mice that lost weight through either bariatric surgical intervention with VSG or weight matched controls were effective at blunting breast cancer progression and reducing tumor burden. However, bariatric surgery only partially reduced obesity accelerated breast cancer progression while weight matched controls effectively blunted growth to a lean-like phenotype.

In patients, weight loss has been shown to improve prognosis after breast cancer has already been diagnosed [13, 45, 46]. Murine models examining weight loss through diet switch, caloric restriction, or time restricted feeding (fasting) support that weight loss impairs tumor progression [21, 47-50]. However, dietary weight loss alone is minimally effective for patients and difficult to maintain. The use of bariatric surgical approaches to induce durable weight loss is increasing in prevalence. In several retrospective studies, bariatric surgery in patients appears to be beneficial to reduce breast cancer risk [23, 51].

Some mechanisms linking obesity-driven breast cancer include elevated adipokines, chronic inflammation, and dampened anti-tumor immune response [33, 52]. We examined multiple factors associated with obesity and metabolic dysfunction, including extent of weight loss, adiposity, mammary fat pad adipocyte size, and local or circulating leptin levels; none were associated with changes in tumor burden in formerly obese mice. However, RNA-seq analysis of the tumor and mammary fat pad demonstrated critical inflammatory pathways regulated by obesity and weight loss. Despite a significant reduction in tumor burden compared to obese HFD-Sham mice, VSG-treated mice demonstrated upregulated mammary fat pad inflammation to levels greater than those of obese mice. Our finding of elevated inflammation in the mammary fat pad after VSG is consistent with several studies reporting inflammation in adipose depots following bariatric surgery in murine models [34-37, 53]. The persistent inflammation identified after bariatric surgery in adipose tissue could be due to adipose remodeling following rapid weight loss, or wound repair signaling from the surgical injury itself. However, these inflammatory changes to the mammary fat pad were uniquely induced by the VSG bariatric surgery, not likely due to surgery itself, since all other groups received a sham surgery as controls. In addition to the mammary fat pad, we report activation of inflammatory and hypoxic pathways in the tumors of mice after VSG but not in other interventions. Therefore, future studies to determine the extent and timing of bariatric surgery associated remodeling in both murine models and humans are warranted. While the murine model presented herein demonstrated successfully stabilized weight loss, most other reports demonstrate weight rebound within a few weeks post-surgery which should be optimized in future cancer studies [29, 30].

We posited that inflammation in the surrounding adipose and tumor led to dramatic elevations in PD-L1 expression and PD-L1+ non-immune cells detected uniquely after VSG [39]. Depressed CD3+ and CD8+ T cell content and dampened expression of T cell cytolytic markers detected in tumors after VSG intervention could have hindered effective anti-tumor immunity after bariatric surgery-associated weight loss. These changes in PD-L1, T cell content and signaling, or cytolytic pathway were not present in the weight matched controls despite this group losing the same amount of weight as VSG intervention. In fact, weight matched controls had significantly elevated cytotoxic T cell tumor infiltration and evidence of cytolytic function which associated with reduced tumor burden.

Tumor inflammation and hypoxia increase expression of PD-L1 within the tumor microenvironment [39]. Inflammation in the obese TME further exacerbates immune checkpoint expression and PD-L1+ cells thus enabling worsened outcomes [42, 54-56]. Patient tumors with high PD-L1 expression are enriched in inflammation, cell adhesion, and angiogenesis pathways [57, 58], which were pathways upregulated in tumors after VSG. Furthermore, tumors from mice that received VSG had high expression of genes that are also enriched in patient tumors that are positive for PD-L1 including *Mefv, Selp, Sema7a*, and *Cysltr1* [58] which are critically linked to responsiveness to ICB. Increasing evidence supports that obesity improves immunotherapy efficacy in melanoma and other cancers and studies in breast cancer are ongoing [59–61]. Here, we report for the first time that anti-PD-L1 was most effective in reducing tumor burden in the mice that received VSG to induce weight loss with restored expression of cytolytic genes. Taken together, we have identified unique anti-tumor efficacy of anti-PD-L1 in mice after VSG.

Next, we determined genes associated with weight loss after bariatric surgery conserved across species. We took advantage of published transcriptomes of subcutaneous adipose tissue from female patients before and after bariatric surgery in comparison with mammary fat pad expression from obese and formerly obese mice after VSG bariatric surgery. We identified a novel weight loss signature specific to bariatric surgery conserved between mice and humans, termed BSAS. Pathways associated with metabolism, remodeling, and immune cells were identified from conserved genes. Because our study consisted of surgical vs dietary interventions and cancer progression, we are in the unique position to compare BSAS transcriptomic changes to tumor outcomes, which we termed T-BSAS. We demonstrate that a subset of 11 key genes in the T-BSAS signature were associated with tumor outcomes in our mouse models. For example, *Ido1*, indoleamine 2, 3-dioxygenase, is part of the rate limiting enzyme that metabolizes L-tryptophan to N-formylkynurenine. The conserved BSAS gene list demonstrated that compared to obese state, *Ido1* is increased by bariatric surgery in both mouse and human. Of note, *Ido1* was not elevated by WM-induced weight loss in our study (data not shown), which suggests that changes in *Ido1* expression could be a specific response to surgically induced weight loss. Over-expression of IDO depletes tryptophan, leading to accumulation of tryptophan metabolites which can induce immunosuppression. Thus, IDO plays a central role in immune escape through reduced CD8+ T cell activation and increased T cell death [62] with multiple IDO inhibitors under investigation [63]. We previously reported that *Ido1* expression in the tumor adjacent mammary fat pad was decreased after anti-PD-1 immunotherapy in obese mice [43]. Thus, the aberrant upregulation of IDO after bariatric surgery-induced weight loss is one potential mechanism limiting anti-tumor immunity in our VSG model that remains under investigation. One limitation of our study is that this study examines just a single syngeneic orthotopically transplanted model wherein we have examined impact of obesity and weight loss on tumor progression and response to immunotherapy. Future work will investigate other cancer models. Additionally, variables such as duration of obesity, extent of surgery, and time post recovery will likely impact immune parameters and should be investigated in pre-clinical and patient settings.

In sum, despite successful and sustained weight loss, tumors in formerly obese mice that received VSG bariatric surgery failed to display sufficiently improved anti-tumor immunity like controls that lost similar amounts of weight. Elevated inflammation in the mammary fat pad and tumor reduced cytotoxic T cells suggested an ineffective anti-tumor milieu after VSG. Anti-PD-L1 immunotherapy was able to improve tumor outcomes in surgical weight loss mice. Ultimately bariatric surgery is the most effective long-term weight loss solution and could be considered in cancer prevention for high-risk obese patients to reduce cancer risk or recurrence. Clinical trials are underway in some severely obese patients with studies examining changes in breast density and breast cancer risk after bariatric surgery [64], reviewed by Bohm *et al* [23]. Understanding how obesity impacts breast cancer anti-tumor immunity and determining effective weight loss strategies to maximize response to therapies will be valuable. In this study, we queried response to ICB in obese and weight loss models, but response to chemotherapy and radiation therapy and combined therapies are also important areas of investigation to advance the field. Because one-third of Americans are considered obese and 9.2% currently severely obese [65], this study is an important first step in understanding bariatric surgery impacts on cancer progression and immunotherapy.

## Methods

### Reagents

All reagents were obtained from Sigma-Aldrich (St. Louis, MO) unless otherwise noted. Fetal bovine serum (FBS, Gibco, Waltham, MA), RPMI 1640 (Corning, Tewksbury, MA), 100X L-glutamine, 100X penicillin/streptomycin HyClone (Pittsburgh, PA), and Gibco 100X antibiotic mix were obtained from Thermo Fisher (Waltham, MA). Matrigel is from (Corning, Tewksbury, MA). Antibodies, compensation beads, and reagents for flow are described in **Table S4** and purchased from (Tonbo, San Diego, CA), Thermo Fisher, and Biolegend (San Diego, CA).

### Mice and diets

Animal studies were performed with approval and in accordance with the guidelines of the Institutional Animal Care and Use Committee (IACUC) at the University of Tennessee Health Science Center and in accordance with the National Institutes of Health Guide for the Care and Use of Laboratory Animals. All animals were housed in a temperature-controlled facility with a 12-h light/dark cycle and *ad libitium* access to food and water, except where indicated. Three-week-old female C57BL/6J (Jackson stock number: 000664) mice were shipped to UTHSC and acclimated 1 week. Four-week-old mice were randomized to either obesogenic high fat diet (HFD, D12492i - 60% kcal derived from fat) or low fat diet (LFD, D12450Ji-10% kcal derived from fat) from Research Diets Inc. (New Brunswick, NJ) for 16 weeks (age 4 weeks to 20 weeks old, study design **Fig 1A**). Mice resistant to diet induced obesity (DIO), as defined by less than 28 grams after 16 weeks of HFD, were excluded from the study. DIO mice received either a bariatric surgery or sham control surgery and dietary intervention as described below.

### Body weight and composition

Body weight was measured 2x/week. Body composition including lean mass, fat mass, free water content, and total water content of non-anesthetized mice was measured weekly using EchoMRI-100 quantitative magnetic resonance whole body composition analyzer (Echo Medical Systems, Houston, TX).

### Vertical Sleeve Gastrectomy

To reduce bariatric surgery-associated weight loss, peri-operative measures included providing liquid diet (Ensure^®^ Original Milk Chocolate Nutrition Shake, Abbott, Chicago, IL) and DietGel recovery (Clear H2O, Portland, ME, ID# 72-06-5022) one day before surgery to all mice. Four hours before surgery, solid food was removed to reduce stomach contents. For 4 hours pre-surgery, mice were maintained half on half off a heat pad in clean new cages. Surgery was performed under isoflurane anesthesia. Vertical sleeve gastrectomy (VSG) was performed as previously described [28] with additional control dietary intervention for comparison of weight loss approaches. The stomach was clamped and the lateral 80% of the stomach was removed with scissors. The remaining stomach was sutured with 8-0 to create a tubular gastric sleeve. All treatment groups not receiving VSG had a sham surgery performed. For sham, an abdominal laparotomy was performed with exteriorization of the stomach. Light pressure with forceps was applied to the exteriorized stomach. For both VSG and sham surgeries, the abdominal wall was closed with 6-0 sutures and skin closed with staples. Mice received carprofen (5mg/kg, subcutaneous, once daily) as an analgesic immediately prior to and once daily for 3 days following surgery. Mice were given 1ml saline at time of surgery. Perioperative procedures were performed in accordance with the literature [66, 67]. For 12 hours post-surgery, mice were maintained half on half off a recovery heat pad. Mice were provided Ensure® liquid diet (as above), DietGel recovery, and solid food pellets *ad libitum* for 48 hours post-surgery. HFD-fed DIO mice receiving VSG (“HFD-VSG”) were maintained on the same HFD for 5 weeks following surgery until euthanasia at study endpoint (**Fig 1A**). Control groups that were lean (“LFD-Sham”) or DIO (“HFD-Sham”) were maintained on respective LFD or HFD diets following sham surgery. For dietary intervention weight loss, DIO mice received sham surgery and were subjected to weight loss intervention following sham surgery for 5 weeks until endpoint. “Weight Matched” (WM) mice were controls to the HFD-VSG mice by weight matching through restricting intake of HFD [68]. On average, mice consumed 1.7g (ranging from 1.0-2.5 g or 8.84 kcal (5.2-13.0 kCal) per day of HFD. Mice were fed at the start of the dark cycle. 78.9% of VSG mice survived to endpoint (30/38).

### Tumor cell implantation

E0771 murine adenocarcinoma breast cancer cell line was originally isolated from a spontaneous tumor from C57BL/6 mouse. E0771-luciferase (luc) is a kind gift from Dr. Hasan Korkaya, Augusta University [43]. Cells were cultured as described previously [43]. Briefly, cells were cultured in RPMI containing 10% FBS, 100 UI/mL of penicillin, and 100 μg/ml streptomycin in a humidified chamber at 37°C under 5% CO_2_. E0771 cells were injected in the left fourth mammary fat pad of 22-week-old C57BL/6J females at 250,000 cells in 100μl of 75% RPMI / 25% Matrigel. When tumors became palpable (typically one week after implantation), tumor growth was monitored 2x/week by measuring the length and width of the tumor using digital calipers. Tumor volume was calculated using the following formula: Volume = (width)^2^ × (length)/2 [43]. No tumors failed to take, and tumor regression was not detected. At the endpoint on day 21 after tumor cell injection, excised tumor mass was determined.

### Immune checkpoint blockade

In a separate experimental cohort limited to HFD-VSG and controls including LFD-Sham, HFD-Sham, and WM-Sham, mice were subjected to the same dietary and surgical study design above (**Fig 1A**). After 20 weeks on LFD or HFD, 24-week-old mice received either a sham or VSG surgery. Two weeks following surgery, mice were injected with E0771-luc cells as above. Immune checkpoint blockade (ICB) included anti PD-L1 antibody (Clone 10F.9G2, #BE0101) and IgG2b isotype control (Clone LTF-2, #BE0090), purchased from BioXcell (West Lebanon, NH). Antibody administration by intraperitoneal (i.p.) injection began three days after E0771 cell injection when tumors were palpable (width of >2.5mm). Mice were injected every third day for 21 days until endpoint (8mg/kg) [69].

### Tissue and blood collection

Three weeks after tumor implantation (i.e., five weeks after surgery), mice were fasted for 4 h and anesthetized. Blood was collected via cardiac puncture into EDTA-coated vials. Plasma was separated from other blood components by centrifugation at 1200×g for 45 min at 12°C. Mammary tumors, tumor adjacent mammary fat pad, unaffected inguinal mammary fat pad, and gonadal adipose were weighed and either flash frozen in liquid nitrogen, placed into a cassette and formalin-fixed, or digested into a single cell suspension for flow cytometry. All frozen samples were stored at −80°C until analyzed.

### Plasma Adipokines

Plasma collected at sacrifice was used for measuring leptin using the Milliplex MAP Mouse Metabolic Hormone Magnetic Bead Panel in the Luminex MAGPIX system (EMD Millipore, Billerica, MA).

### Flow cytometric analysis of tumors and adjacent mammary adipose tissue

Flow cytometry analysis was done as previously described [43]. In brief, excised tumors (200 mg) were dissociated in RPMI media containing enzyme cocktail mix from the mouse tumor dissociation kit (Miltenyi Biotec, Auburn, CA) and placed into gentleMACS dissociators per manufacturer’s instructions. Spleen single cell suspensions were obtained by grinding spleens against 70μm filter using a syringe plunger. Following red blood cell lysis (Millipore Sigma, St. Louis, MO), viability was determined by staining with Ghost dye (Tonbo Biosciences Inc.) followed by FcR-blocking (Tonbo). Antibodies in **Table S4** were titrated, and separation index was calculated using FlowJo v. 10 software. Cells were stained with fluorescently labeled antibodies and fixed in Perm/fix buffer (Tonbo). Stained cells were analyzed using Bio-Rad ZE5 flow cytometer. Fluorescence minus one (FMO) stained cells and single color Ultracomp Beads (Invitrogen, Carlsbad CA) were used as negative and positive controls, respectively. Data were analyzed using FlowJo v 10 software (Treestar, Woodburn, OR). Total immune cells from tumor and tumor adjacent mammary fat pad (including tumor draining lymph node, TdLN) were gated by plotting forward scatter area versus side scatter area, single cells by plotting side scatter height versus side scatter area, live cells by plotting side scatter area versus Ghost viability dye, and immune cells by plotting CD45 versus Ghost viability dye. T cells were gated as follows in tumor CD3+ T cells (CD3+), and CD8+ T cells (CD3+, CD8+). Non-immune cells were gated as CD45- and mean fluorescent intensity (MFI) for PD-L1. Gates were defined by FMO stained controls and verified by back-gating of cell populations. Gating schema is shown in supplemental figure 3 **(Fig. S3)**.

### RNA sequencing (RNA-seq)

mRNA was extracted from tumor tissue using RNeasy mini kit (QIAGEN, Germantown, MD) and mammary fat pad tissue using a kit specific for lipid rich tissue (Norgen Biotek, Ontario, Canada). The integrity of RNA was assessed using Agilent Bioanalyzer and samples with RIN >8.0 were used. Libraries were constructed using NEBNext® Ultra™ RNA Library Prep Kits (non-directional) for Illumina, following manufacturer protocols. mRNA was enriched using oligo-dT beads. Libraries were sequenced on NovaSeq 6000 using paired-end 150 bp reads. There was no PhiX spikein. Data was analyzed as described previously [43, 70]. RNA-seq statistical differences between experimental groups were determined as described previously [43]. In brief, Benjamini-Hochberg procedure was used to control false discovery rate (FDR) for adjusted P value. RNA-seq data has been uploaded as GEO GSE174760, GSE174761, and GSE174762. Transcript-level abundance was imported into gene-level abundance with the R package tximport. Genes with low expression were identified and filtered out from further analysis using filterByExpr function of the edgeR package in R software. Voom transformation function was applied to normalize log2-cpm values using mean-variance trend in the limma software package. ClaNC was used to create classifier genes that characterize the groups of interest for semi-supervised heatmaps. Database for Annotation, Visualization and Integrated Discovery (DAVID) v6.8 was used for pathway analysis [71]. Immune infiltration estimations based on bulk gene expression data from RNA-seq was plotted using TIMER2.0 [72] and cell-type identification estimating relative subsets of RNA transcripts (CIBERSORT) [73].

### Bariatric Surgery Patient RNA-seq

Patient gene expression from subcutaneous adipose tissue pre- and post-bariatric surgery was downloaded from GSE65540 [37] and counts were normalized using counts per million (CPM). EdgeR was used for differential expression analysis and significance was defined as adjusted p-value of < 0.1. Benjamini-Hochberg was used to calculate the FDR. Mouse and human Venn diagram was created using the interactive Venn website.

### Gene expression

Total RNA was isolated from tumors and reversed transcribed to cDNA using High-Capacity cDNA Reverse Transcription Kit (Applied Biosystems). qRT-PCR was performed with iTaq Universal SYBR Green Supermix (Bio-Rad). Primers span an exon-exon junction and were designed with Primer-BLAST (NCBI). Relative gene expression was calculated normalized to 18S transcript with 2^-ΔΔCt^. Primer sequences are:

*Ifng* F:GGATGCATTCATGAGTATTGC, *Ifng* R:GTGGACCACTCGGATGAG,
*Prf1* F:GAGAAGACCTATCAGGACCA, *Prf1* R:AGCCTGTGGTAAGCATG,
*Gzmb* F:CCTCCTGCTACTGCTGAC, *Gzmb* R:GTCAGCACAAAGTCCTCTC,
*18S* F: TTCGGAACTGAGGCCATGATT, *18S* R:TTTCGCTCTGGTCCGTCTTG

### Histology and quantification

Tumors and normal 4^th^ mammary fat pads, (contralateral to the injected tumor bearing mammary fat pad) were isolated at the time of sacrifice and fixed in 10% formalin. Formalin fixed paraffin embedded (FFPE) sections from tumors and adipose were cut at 5 μm thickness. FFPE sections were stained with Hematoxylin and Eosin and-scanned by Thermo Fisher (Panoramic 250 Flash III, Thermo Fisher, Tewksbury, MA) scanner and adipocyte area of N=50 adipocytes were quantified using software (Case Viewer) along the longest diameter per adipocyte.

### Statistics

Statistical differences between experimental groups were determined using One-way or Two-way ANOVA (as noted in figure legends) with Fisher’s LSD test for individual comparisons. Outliers were identified and excluded based on the ROUT method with Q=1%. For body weight, body composition, and tumor volume over time within animals, data was treated as repeated measures. All statistics were performed using statistical software within Graphpad Prism (Graphpad Software, Inc., La Jolla CA). All data are shown as mean ± standard error of the mean (SEM). P values less than 0.05 were considered statistically significant.

### Study approval

Animal studies were performed with approval and in accordance with the guidelines of the Institutional Animal Care and Use Committee (IACUC) at the University of Tennessee Health Science Center and in accordance with the National Institutes of Health Guide for the Care and Use of Laboratory Animals.

## Author contributions

Conceptualization: LMS, JFP, LM
Funding acquisition: LMS, DNH, JFP, LM
Formal analysis: LMS, MC, HJ, UT, RS
Investigation: LMS, MC, EBK, MCL, BRC, JCC, NAJ,
Methodology: LMS, MC, DD, TNM, AKP, BT, JAC, JFP, LM
Project administration: LMS, LM
Resources: JAC, DNH, JFP, LM
Supervision: JAC, MJD, DNH, JFP, LM
Writing – original draft: LMS, LM
Writing – review & editing: LMS, LM, MJD, JFP, DNH, JAC

We acknowledge support from the following funding sources:

National Institutes of Health grant NCI R01CA253329 (LM, JFP, MJD)
National Institutes of Health grant NCI R37CA226969 (LM, DNH)
The Mary Kay Foundation (LM)
V Foundation (LM, DNH)
National Institutes of Health grant NIDDK R01DK127209 (JFP)
Tennessee Governor Pediatric Recruitment Grant (JFP)
Tennessee Clinical and Translational Science Institute (JFP)
Transdisciplinary Research on Energetics and Cancer R25CA203650 (LMS)
American Association for Cancer Research Triple Negative Breast Cancer Foundation Research Fellowship (LMS)
National Institutes of Health grant NCI F32 CA250192 (LMS)
The Obesity Society/Susan G. Komen Cancer Challenge award 2018 (LMS)
National Institute of Health grant NCI U24CA210988 (DNH)
National Institute of Health grant NCI UG1CA233333 (DNH)
National Institute of Health grant NCI R01CA121249 (JAC)
UT/West Cancer Research Institute Fellowship 2019 (JCC)
NIH Medical Student Research Fellowship (MSRF) Program 2019 (NAJ)

We thank Daniel Johnson from UTHSC Molecular Resource Center.

## Data Availability Statement

The data generated in this study are available within the article and its supplementary data files.

The RNA-seq data generated in this study are publicly available in NCBI GEO GSE174760 of tumor RNA-seq and NCBI GEO GSE174761 of mammary fat pad RNA-seq.

