## Supplemental Tables and Figures for "Response to immune checkpoint blockade improved in pre-clinical model of breast cancer after bariatric surgery"

**Supplemental Table 1. Multiple comparisons of body weight after surgery over time.**
*p<0.05, **p<0.01, ***p<0.001, ****p<0.0001. Two-Way ANOVA with Fisher's LSD test.
Low fat diet (LFD), High fat diet (HFD), Vertical sleeve gastrectomy (VSG), Weight-Matched (WM)

| | Weeks after surgery | | | | | | | | | | |
|---|---|---|---|---|---|---|---|---|---|---|---|
| | 0 | 0.5 | 1 | 1.5 | 2 | 2.5 | 3 | 3.5 | 4 | 4.5 | 5 |
| LFD-Sham vs. HFD-Sham | **** | **** | **** | **** | **** | **** | **** | **** | **** | **** | **** |
| LFD-Sham vs. HFD-VSG | **** | **** | **** | **** | *** | *** | ** | *** | ** | *** | **** |
| LFD-Sham vs. WM-Sham | **** | **** | **** | **** | *** | ** | ** | ** | * | * | ** |
| HFD-Sham vs. HFD-VSG | ns | ns | *** | **** | **** | **** | **** | **** | **** | **** | **** |
| HFD-Sham vs. WM-Sham | ns | ns | * | ** | **** | **** | **** | **** | **** | **** | **** |
| HFD-VSG vs. WM-Sham | ns | ns | ns | ns | ns | ns | ns | ns | ns | ns | ns |

**Supplemental Table 2. Multiple comparisons of tumor volume over time.**

*p<0.05, **p<0.01, ***p<0.001, ****p<0.0001. Two-Way ANOVA with Fisher's LSD test.
Low fat diet (LFD), High fat diet (HFD), Vertical sleeve gastrectomy (VSG), Weight-Matched (WM)

| | Weeks after tumor implantation | | | | |
|---|---|---|---|---|---|
| | 1 | 1.5 | 2 | 2.5 | 3 |
| LFD-Sham vs. HFD-Sham | *** | ** | **** | *** | **** |
| LFD-Sham vs. HFD-VSG | * | ns | ns | * | ** |
| LFD-Sham vs. WM-Sham | ns | ns | ns | ns | ns |
| HFD-Sham vs. HFD-VSG | ns | * | * | ns | * |
| HFD-Sham vs. WM-Sham | ** | ** | **** | *** | **** |
| HFD-VSG vs. WM-Sham | ns | ns | ns | ** | ** |

**Supplemental Table 3. Conserved differentially expressed genes in subcutaneous adipose/mammary fat pad in obese and bariatric surgery patients and mice.**

| Gene symbol |
|---|
| Ret |
| Ddah1 |
| Hp |
| Lpgat1 |
| Nek6 |
| Ankrd50 |
| Sparc |
| Tuft1 |
| Rab20 |
| Chka |
| Dgki |
| Lep |
| Tgm1 |
| Itga1 |
| Tmem125 |
| Cd200 |
| Slc7a4 |
| Msc |
| Usf2 |
| Ephb3 |
| Cntn2 |
| Lgals7 |
| Mast4 |
| Tusc1 |
| Aldoc |
| Klhl5 |
| Arhgap20 |
| Setd7 |
| Thoc2 |
| Nap1l1 |
| Nkiras1 |
| Cmtm8 |
| Serpinf1 |
| Psme4 |
| Col4a1 |
| Clca2 |
| Nrp2 |
| Ficd |
| Kctd10 |
| Rtn4rl1 |
| Eif4b |
| Vgll3 |
| Slc15a4 |
| Slc35g1 |
| Pde8a |
| Mid1 |
| Tarsl2 |
| Sema3c |
| Pcdh7 |
| Vps13a |
| Amn1 |
| Ido1 |
| Npr3 |
| Srsf4 |

**Supplemental Table 4. Antibodies, fluorophores, and companies**.

| **Antibody** | **Fluorophore** | **Company** | **Dilution Factor** | **Catalog number** |
|---|---|---|---|---|
| Anti-Mouse CD45 | violetFluor 450 | Tonbo Biosciences | 0.025 | 75-0451-U025 |
| Anti-Mouse CD3ε | Brilliant Violet 785 | BioLegend | 0.025 | 117339 |
| Anti-Mouse CD8a | FITC | Tonbo Biosciences | 0.0104 | 35-0081-U025 |
| Anti-Mouse CD274 | Brilliant Violet 711 | BioLegend | 0.10 | 124319 |
| Anti-Mouse PD-1 | Brilliant Violet 421 | Biolegend | 0.10 | 135217 |
| **Reagents** | | | | |
| UltraComp eBeads™ | | Thermo Fisher | | 01-2222-41 |
| ArC™ Amine Reactive Compensation Bead Kit | | Thermo Fisher | | A10628 |

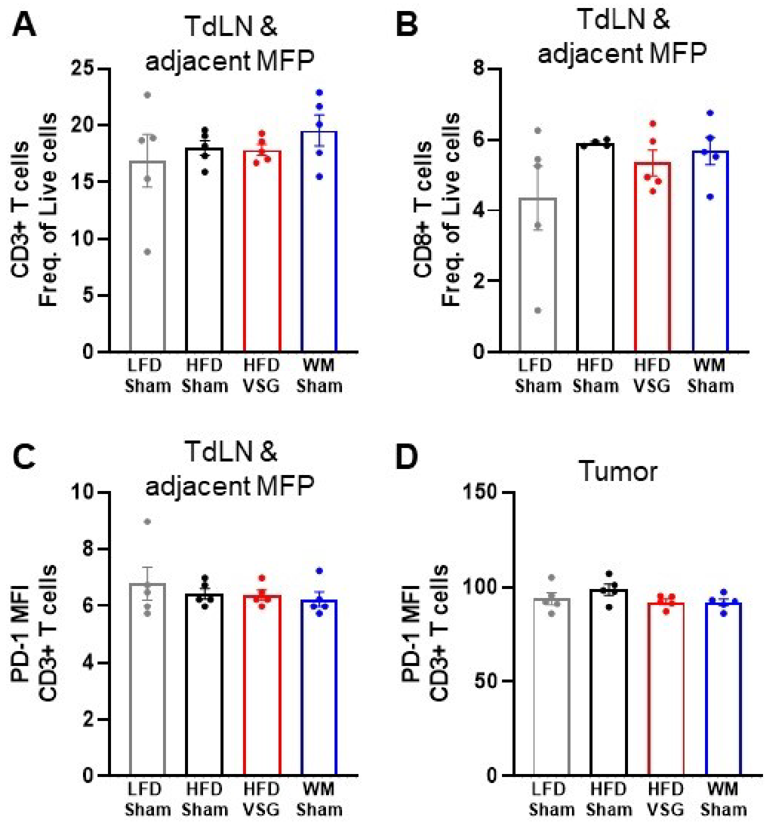

**Supplementary Figure 1. Tumor draining lymph node and tumor infiltrating CD3+ and CD8+ T cell frequencies were not changed, nor was CD3+ PD-1 expression.**

Flow cytometric analysis of tumor draining lymph node (TdLN) and tumor adjacent mammary fat pad (MFP) tissue (**A**) CD3+ T cells and (**B**) CD8+ T cells are shown as frequency of total live cells. Mean fluorescent intensity (MFI) of PD-1 on CD3+ T cells in (**C**) TdLN and tumor adjacent MFP and in (**D**) tumor is shown. (A-D) Mean ± SEM N=5. One-way ANOVA with Fisher's LSD test.

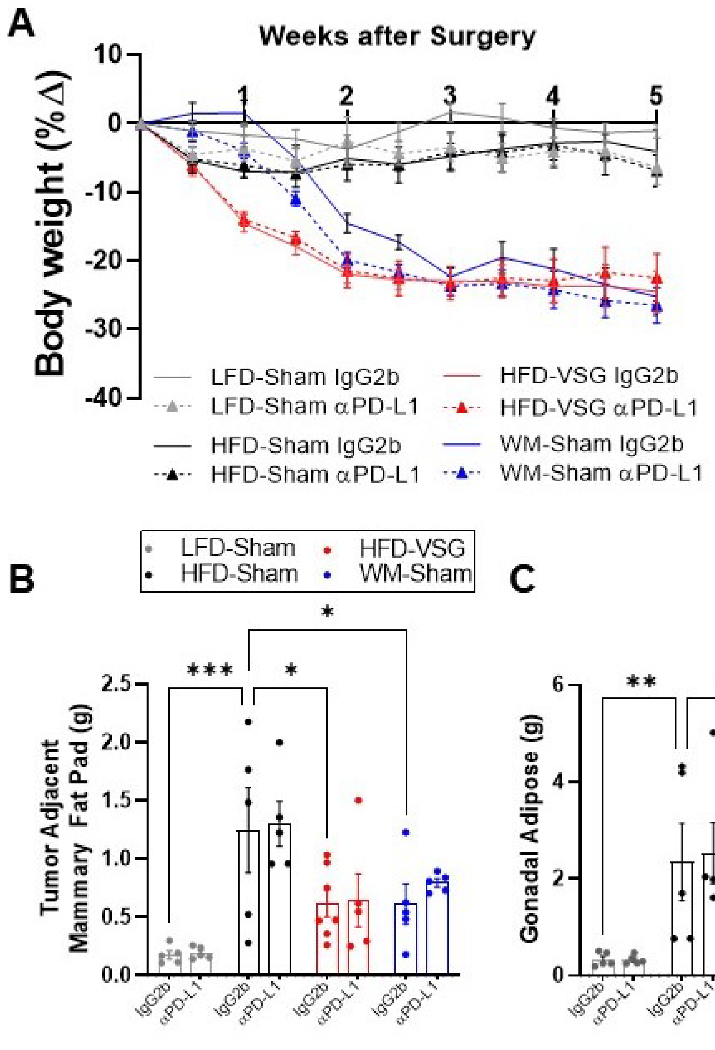


**Supplementary Figure 2. Immune checkpoint blockade did not alter body weight or adiposity.**

(**A**) Percent body weight change in mice after weight-loss interventions is reported until endpoint. (**B**) Tumor adjacent mammary fat pad and (**C**) gonadal adipose weight at endpoint is reported. Mean ± SEM. N=5-8. Two-way ANOVA with Fisher's LSD test. *p<0.05, **p<0.01, ***p<0.001.

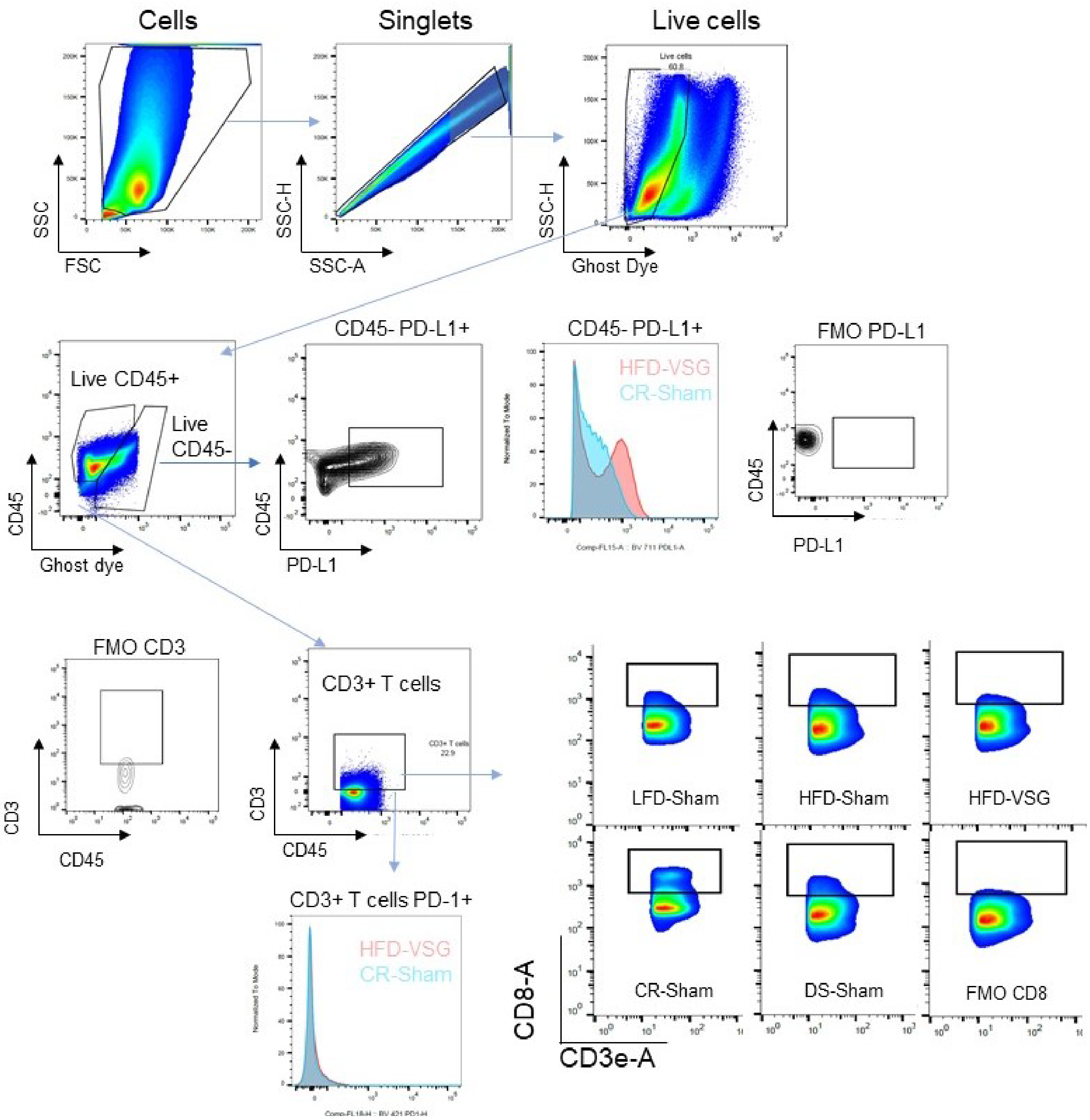


**Supplemental Figure 3. Gating schema for flow cytometric analysis of immune cells in tumor single cell suspensions.**

Total cells from tumor or tumor adjacent mammary fat pad (including tumor draining lymph node, TdLN) were gated by plotting forward scatter area versus side scatter area, single cells by plotting side scatter height versus side scatter area, live cells by plotting side scatter area versus Ghost viability dye, and immune cells by plotting CD45 versus Ghost viability dye. T-cells were gated as follows in tumor CD3+ T cells (CD3+), and CD8+ T cells (CD3+, CD8+). MFI of PD-1 was measured in CD3+ PD-1+ cells. Non-immune cells were gated as CD45-, PD-L1+, and MFI for PD-L1.
